# Cell-to-cell variability and gain of methylation at polycomb CpG islands as a hallmark of aging

**DOI:** 10.64898/2026.03.09.710505

**Authors:** Hagit Masika, Shmuel Ruppo, Stephen J. Clark, Marc Jan Bonder, Ferdinand von Meyenn, Merav Hecht, Shari Orlanski, Efrat Katsman, Oriya Vardi, Abraham Zlotogorski, Sharona Elgavish, Yuval Dor, Wolf Reik, Tommy Kaplan, Howard Cedar

## Abstract

Aging is a complex multifactorial process that affects cellular function and tissue homeostasis over time. Despite extensive research, the molecular mechanisms driving cellular aging remain poorly understood^1,2^. Many studies have focused on changes in DNA methylation as an indicator of aging^3^. In particular, the degree of methylation at polycomb CpG islands has been shown to be predictive of phenotypic changes associated with aging^4,5^. Since many age-related pathological processes, are thought to be of single-cell origin (e.g. cancer), we questioned whether polycomb DNA methylation also occurs preferentially in a subset of cells within the overall population. Using single-cell whole-genome methylation data from multiple ages and tissues, we identify Average Polycomb CpG Methylation as a hallmark of cellular aging. This revealed that aging occurs at varying rates within specific cells, with faster proliferating cells showing accelerated levels. Gene expression analysis in “young” and “old” single cells identified changes in immune response, translation regulation, tumorigenesis, neurodegeneration and other cellular processes associated with aging. These results challenge traditional models of homogeneous cellular aging and suggest that aging itself is a highly individualized process at the single-cell level that may be driven by programmed changes in polycomb CpG island DNA methylation.

## Introduction

DNA methylation represents a major epigenetic marker that plays a role in gene regulation throughout development and its profile in the genome is largely responsible for determining stable cell-type identity^6^. In keeping with this important concept, the DNA methylation landscape has also emerged as a fairly accurate and reliable index of chronological age^3^.

On the basis of early studies on the changes in DNA methylation occurring in cancer, it was shown that many of these alterations are identical to dynamic events that take place during aging. Furthermore, it appears that this process is not stochastic, but actually involves a form of global programming in the sense that it is specifically on CpG islands whose DNA sequence has the ability to bind the protein complex, polycomb^7^. One of the most outstanding features of this methylation pattern is that this same mark serves as an index of aging in all cell types, each at a different rate. This suggests that polycomb CpG island methylation is a fundamental, developmentally programmed phenomenon that may actually play a causal role in the aging process. As partial proof of this idea, we have demonstrated that the level of polycomb-island methylation in different tissues of the body is actually a reliable predictor of lifetime cancer risk, even though this was measured in normal cells, prior to the appearance of any tumors^5,8^.

Several years ago, our laboratory made the interesting observation that when DNA from normal tissues was subjected to single-molecule bisulfite sequencing, it was found that the DNA methylation content of polycomb CpG islands is skewed, with some molecules being highly methylated, while the large majority have almost no methylation. These results, as well as previous single-cell methylation data showing the presence of “young” cells in chronologically old tissue^9,10^, suggested, but did not prove, that aging-associated DNA methylation may occur preferentially in a small number of individual cells as opposed to the entire population^8^. Taken together with the fact that all tumors are initially generated from single individual cells, we decided to test the idea that aging – at the single-cell level – may occur at different rates among seemingly homogenous cells, adding a cellular aspect to the more commonly perceived population model of organ aging.

In order to test this hypothesis, we initiated a project to measure whole-genome DNA methylation in single cells and then created a new computational strategy to quantitate^5,11^ the degree of aging on a cell-cell basis. As determined from our own experiments, as well as extensive published data for a large variety of different cell types in mouse and human, we demonstrate that the phenomenon of aging as defined by our polycomb CpG island DNA methylation index takes place at highly variant rates in different individual cells that make up the specific cell type. Thus, at any chronological age, each specific tissue or cell type of the body is actually composed of a heterogenous mixture of cells with a relatively low aging index, along with cells that advance more rapidly. Taken together with gene expression analysis, which confirms this concept at the functional level, these studies provide an entirely new perspective on what constitutes physiological aging.

## Results

During early development, almost all gamete-derived DNA methylation is erased^12^. Then, at the time of implantation, the overall genome gets remethylated, but CpG islands are protected from this process and thus remain unmethylated, a pattern that, once formed, is then faithfully maintained in all tissues throughout life^13^ . Nonetheless, a subset of CpG island-regions, those specifically bound by the polycomb complex, have been found to undergo slow de novo methylation as a function of age^14^, and this process is probably mediated by the presence of Ezh2, a subunit of the PRC2 complex, which has been shown to have a binding domain for DNA methylases^15^.

### Polycomb CpG island methylation is an indicator of age

A number of laboratories have analyzed DNA methylation data from different cell types in the body, using machine-learning algorithms to derive indices proportional to chronological age, both in human and mouse^3^. As opposed to this empirical approach, we have introduced the concept of employing the DNA methylation levels at hundreds of polycomb-bound CpG islands, as a general programmed indicator of aging. When this instructive method was applied to published TCGA data from a variety of different normal cell types, we were able to verify that this index is indeed a reliable measure of aging^5^. Our analysis of whole-genome DNA methylation data (Supplementary Fig. 1)^10,16–18^ yielded a similar picture, thereby confirming the idea that this programmed molecular process is closely aligned with aging.

### Experimental strategy

To study the heterogeneity of polycomb CpG island DNA methylation as an index of epigenetic aging, we moved from average methylation in bulk tissues, to a “single-cell” approach. As a first step, we chose to examine individual intestinal crypts. Each crypt contains about 100 cells, all derived from resident stem cells, and it has been demonstrated that this represents a fairly pure single-cell clonal population^19^. Since DNA methylation patterns are faithfully maintained through cell division, each crypt contains cells with identical profiles. Single crypts were isolated and subjected to whole-genome bisulfite sequencing (WGBS). These data were then analyzed to determine their DNA methylation pattern. A major problem in the measurement of whole-genome DNA methylation is its inherent inefficiency caused by the relative scarcity of CpGs in the genome, as well as the limited sequencing coverage per each cell clone. We overcome this by assuming that the gain of de novo methylation occurs stochastically and uniformly across hundreds of polycomb CpG islands (Supplementary Datasets 1,2), and then average their DNA methylation levels^5,11^. After confirming that this prediction is indeed robust (Methods), we calculated the average polycomb CpG island DNA methylation index in each cell clone. This allowed us to obtain accurate values even though most of these specific regions individually were not covered by a significantly sufficient number of reads.

### Methylation aging in single colonic crypts

Using this averaging approach, we then analyzed the results obtained from 87 individual crypts taken from 22 and 28 wk old mice. As shown in Fig. 1a, the mean aging index for the samples taken from the 28 wk animals have a higher range of polycomb CpG island methylation levels as compared to those obtained from 22 wk animals, clearly indicating that average DNA methylation levels in polycomb CpG islands are indeed correlated with chronological age. At the same time, the individual-crypt distribution of this index is significantly skewed (p<0.003, p<1e-5 for 22wk and 28wk, respectively, Shapiro-Wilk test), with some crypts showing methylation levels much higher than what would be expected by random simulations using a stochastic model (Methods).

**Figure 1:**
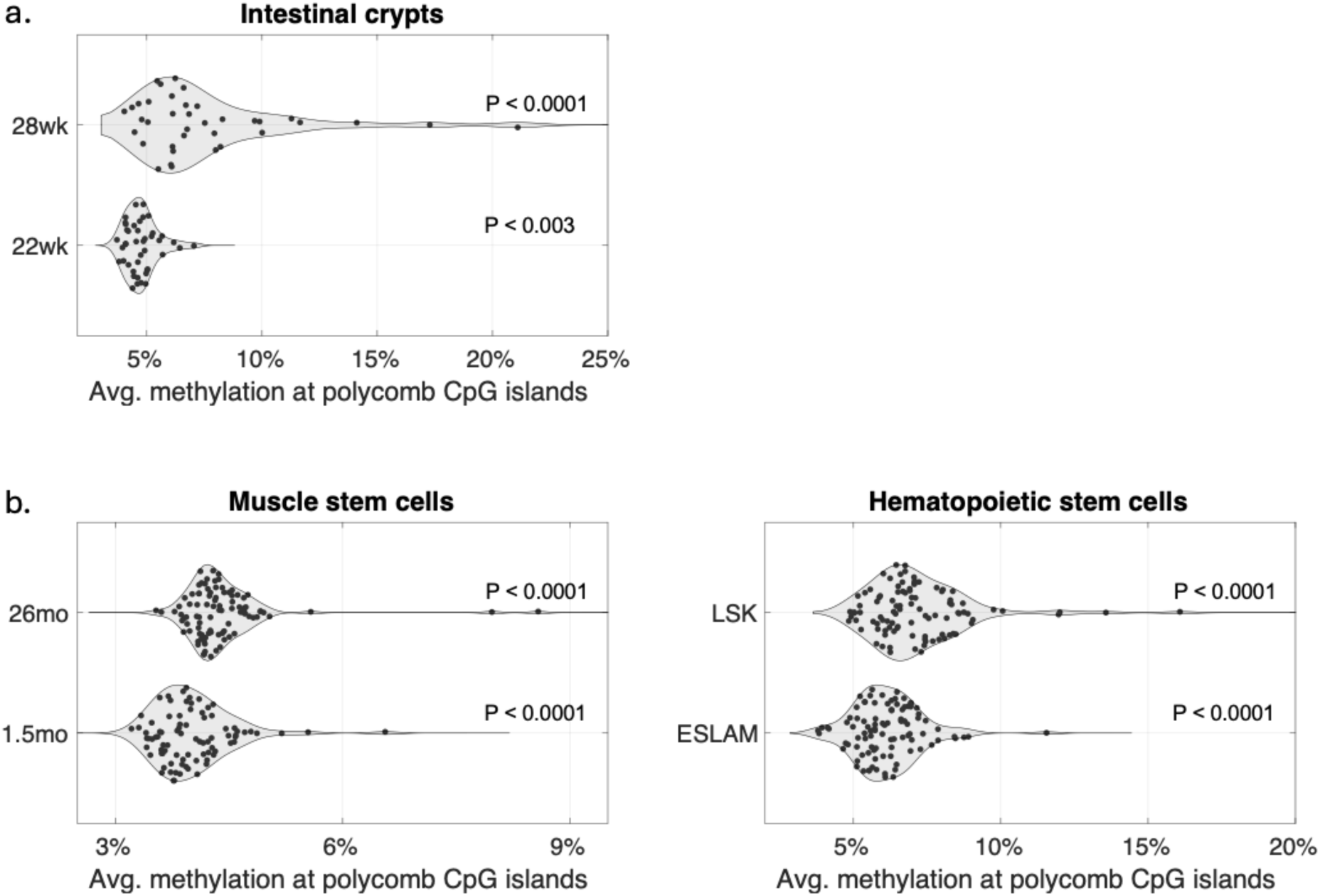
Distribution of average DNA methylation levels across polycomb CpGs islands (n=2,975) in individual crypts / cells from various tissues. (a) Average methylation in intestinal crypts from 22-week-old (n=43) and crypts from 28-week-old mice (n=37), showing age-related gain of methylation. Each dataset was compared to a simulated set of cells, stochastically sampled from the null hypothesis (Methods). It should be noted that the large variation between crypts observed for polycomb CpG methylation does not reflect the overall methylation pattern of every crypt taken from a single mouse (28w), which is highly uniform (Shapiro Wilk, p<0.7), in keeping with the fact that each crypt is monoclonal and derived from a single stem cell. (b) Same as (a), for stem cell populations. Left panel shows the distribution of polycomb methylation in muscle stem cells from young (1.5 months, n=84) and old (26 months, n=94) mice^9^. Right panel: Hematopoietic stem cells (HSCs), including endothelial stem-like cells (ESLAM, n=97) and LSK cells (Lin−Sca1+c-Kit+, n=100), derived from the bone marrow of 8-10-week-old mice^43^.

### Single-cell polycomb CpG island DNA methylation in aging

To test whether the aging dynamics observed in cells of the intestine is typical of other tissues in the body, we next used the same averaging approach to analyze the single-cell methylomes of white blood cells (n=1,132) at ages 10, 36, 77 and 100 weeks^10^. For each individual cell, we calculated an aging index equal to the average methylation at n=2,975 polycomb CpG islands. When graphed, the data show a monotonic increase in polycomb CpG island methylation across the mouse life span (Fig. 2a, left). We also observed a strongly skewed distribution, with the majority of cells exhibiting relatively low levels of methylation, while others appeared to have a much higher index. This is especially prominent in cells from 77- or 100-week mice (p<1e-5, Shapiro-Wilk test). These age distributions differ significantly from the cell-to-cell variability expected using a stochastic simulation of single-cell data assuming a uniform rate (p<1e-5). Instead, this skewed distribution strongly supports the concept that, in addition to a slow homogenous gain of de novo methylation at polycomb CpG islands, some individual cells may age independently at an accelerated rate.

**Figure 2:**
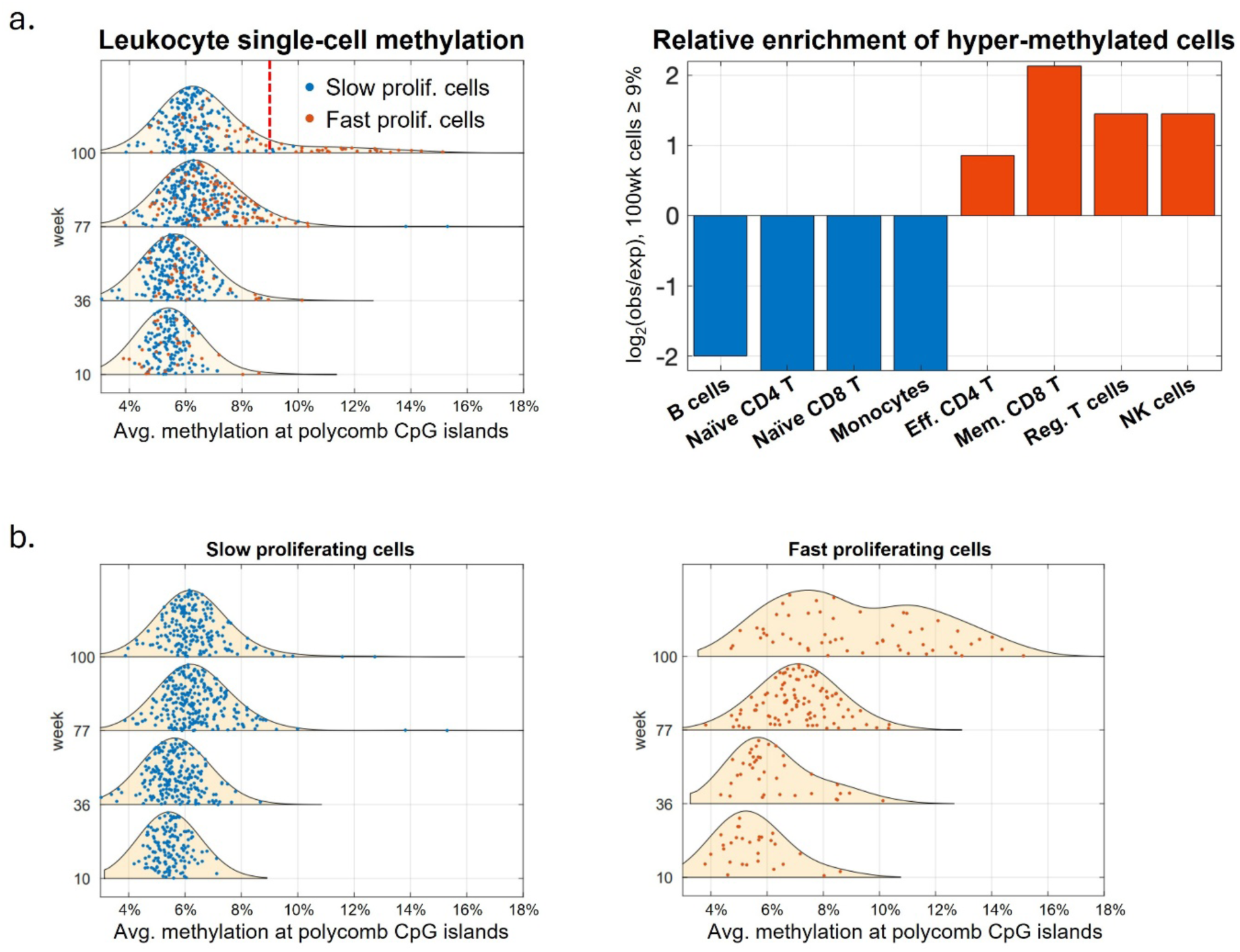
Gain of polycomb CpG island methylation and cell-to-cell variability in white blood cells. **(a)** Left: Ridgeline plots displaying the distribution of polycomb CpG island DNA methylation in n=1,132 white blood cells, at four different chronological ages: 10, 36, 77, and 100 weeks. For each individual cell, the aging index was calculated based on the degree of de novo methylation at a set of 2,975 polycomb CpG islands. Blue dots represent non-proliferating cells, whereas red dots represent proliferating cells. Average methylation levels and standard deviations monotonically increase with time, from 5.5%±0.77% to 7%±2%. Right: Bar plot showing the relative abundance (log2 of observed/expected ratio) of each specific cell type among 100-week old cells with polycomb methylation index ≥ 8.9% (dashed red line). Blue: slow-proliferating cell types (B cells, naïve CD4 T cells, naïve CD8 T cells, monocytes). Red: fast proliferating cells (Effector CD4 T cells, memory CD8 T cells, regulatory T cells, NK cells); **(b)** Ridgeline plots as in (a), for slow-proliferating (blue) and fast proliferating (red) cell populations at each chronological age cell group. Avg. methylation levels (and std) for proliferating cells vary from 5.59%±1.27% for 10 wk up to 9.3%±2.9% for 100 wk.

In order to rule out the possibility that this skewing may be due to the presence of multiple cell types, we used the single-cell transcriptomics data, that was collected for the same individual cells^10^ to infer the blood cell type of each individual single cell. We then examined the relative proportion of each cell type as a function of age-associated hyper-methylation. Specifically, when focusing on week 100 cells, we observed a clear demarcation at an average methylation threshold of 8.9%. Some specific cell types (EffCD4T, MemCD8T, RegT and NK) show a ∼2-4-fold enrichment within the accelerated methylation (rapidly aging) distribution tail, while others (B cell, NveCD4T, NveCD8T, Mono) are in the slow aging group (Fig. 2a). These data immediately suggested that the two groups may correspond to their ability to proliferate, where the accelerated methylation group is associated with higher proliferation rates as opposed to the other cell types that proliferate with slower kinetics. This was then confirmed by analyzing the polycomb CpG island methylation index of slow-proliferating as opposed to fast-proliferating cells (Fig. 2b). The slow-proliferating cells show a consistent increase in polycomb CpG island methylation with a near-normal distribution across individual cells. Conversely, the fast-proliferating cells appear to have a more complex pattern, suggesting a slow gradual increase in methylation for most cells, as well as a growing age-dependent sub-population of cells, showing an accelerated rate of methylation gain. In general, the fast-proliferating cells accumulate polycomb CpG island methylation at a faster rate than slow-proliferating cells (Supplementary Fig. 1).

### Single-cell aging occurs in many cell types

In order to verify our findings in a more general manner, we next examined single-cell DNA methylation in a variety of additional cell types, using data previously reported in the literature (Fig. 1b). Analysis of polycomb CpG island methylation in purified muscle stem cells from relatively young (1.5 month old) animals, for example, showed a skewed distribution from the expected normal (see legend). While most cells show low levels of polycomb CpG island methylation, some outlier cells already show high methylation levels. This skewing was even more pronounced in muscle stem cells derived from 26 mo. old animals (p≤1e-5, Jarque-Bera kurtosis test). A similar pattern was also seen in hematopoietic stem cells isolated from the bone marrow of 8-10 wk mice, where both multipotent hematopoietic cells (LSK) and progenitor cells (ESLAM) show a much wider range of methylation index than expected (Fig. 1b).

The phenomenon of single-cell epigenetic aging was also seen in human cells. Peripheral myeloid and lymphoid progenitor cells (CMP and CLP) taken from 60-65 year olds are not distributed in a normal manner, and the same seems to be true for megakaryocytes taken from the bone marrow of a 50-year old patient. These cells give an illustrative example of the wide range of variability in individual cell methylation, with some cells showing a methylation index of 2% and others 12%. Strikingly, a similar pattern was observed in hematopoietic stem cells (HSCs) taken from either fetal liver or cord blood, very early stages in the human lifetime scale (Fig. 3).

**Figure 3:**
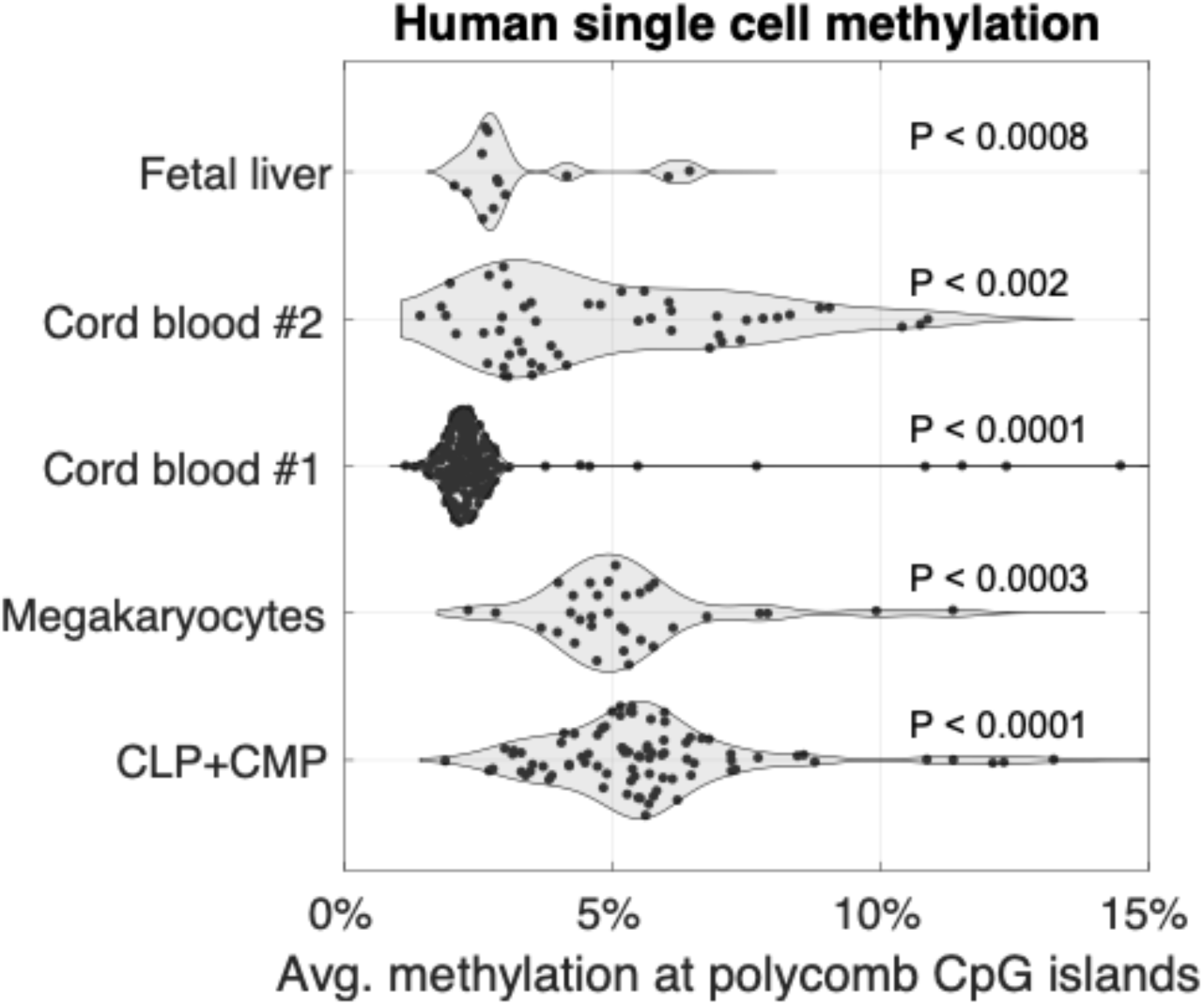
Distribution of average polycomb CpG island methylation in human single cells. Methylation indices were calculated for various single cell populations^44^, using human polycomb CpG islands (n=500) (Supplementary Dataset 2). Data include fetal liver (n=13 cells), two cord blood samples (n=225, n=50), bone marrow megakaryocytes from a 50-year-old patient (n=33), and common lymphoid progenitor (CLP) and common myeloid progenitor (CMP) cells (n=84), from 60-65-year-old patients^43^. Each dataset was compared to a simulated set of cells, stochastically sampled from the null hypothesis (Methods).

### Demethylation in aging

Another relatively universal aspect of aging is the widespread loss of methylation at late-replicating domains, associated with lamin B (LADs) on the nuclear envelope^20,21^. In an inverse manner to de novo methylation of polycomb CpG islands, these sequences have been found to be partially methylated in cancer. In order to determine whether these two separate processes are mechanistically coordinated, we developed a numerical index to quantify the demethylation of the sites that undergo the most pronounced change (Solo-WCGW CpGs)^20^ and applied this to the analysis of single-cell WGBS. We then generated scatter plots to test the relationship between the demethylation and de novo methylation indices on a single-cell basis at all chronological ages. While all lymphocytes appear to undergo CpG island methylation aging at similar rates, there were marked differences in LAD demethylation between each specific cell type (Fig. 4a).

**Figure 4:**
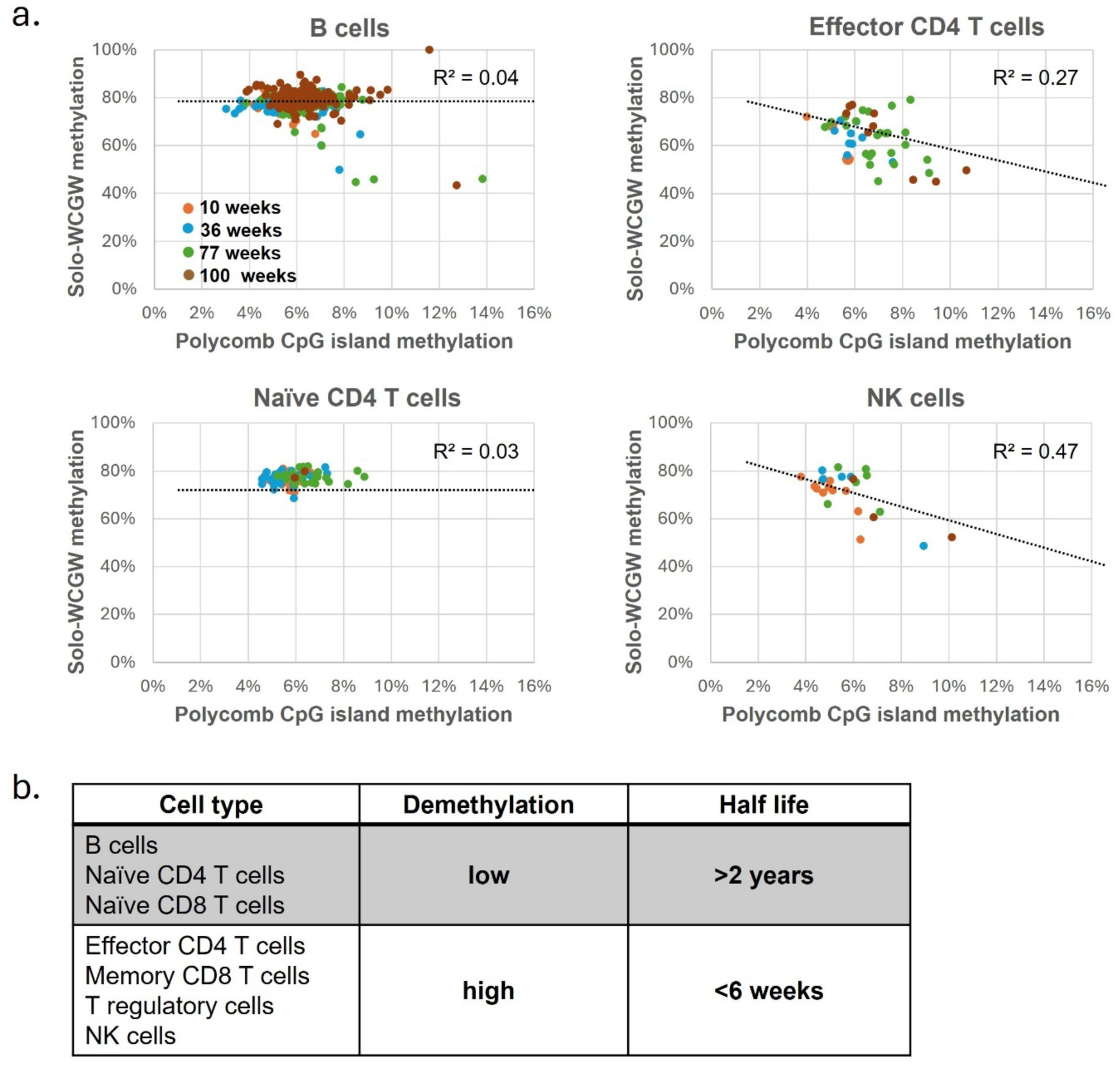
Single-cell comparison of de novo methylation at polycomb CpG islands vs. loss-of-methylation at partially methylated domains (PMDs). **(a)** Scatterplots compare the average methylation across polycomb CpG islands (X-axis) vs average methylation at solo-WCGW CpGs at lamina-associated partially methylated domains (PMDs) Y-axis)^20^. Correlations are shown for four cell types: B cells, effector CD4 T cells, naïve CD4 T cells, and NK cells, at four ages (10, 36, 77, and 100 weeks, color-coded). **(b)** Different white blood cell types are represented according to the degree of demethylation relative to their *in vivo* proliferation rates (B cells^45^, Naïve CD4 T cells, Naïve CD8 T cells^46,47^, Effector CD4 T cells^48^, Memory CD8 T cells^49^, T regulatory cells^50^, NK cells^51^).

B cells, for example, do not seem to undergo any appreciable age-associated demethylation, with single cells from all ages retaining their originally high LAD methylation levels. Naïve CD4 cells behaved in a similar manner. In contrast, NK and Effector CD4 T cells reveal considerable demethylation with aging in a manner that is clearly proportional to age-related de novo methylation, indicating that these processes are in some way linked. The same is true for T-reg and memory cells as well (not shown). As demonstrated previously, the rate of demethylation with aging is highly dependent on DNA replication^20^. In keeping with this observation, analysis of the cell types examined in this study indeed show that the kinetics of age-related demethylation is approximately proportional to their proliferation rate in vivo (Fig. 4b) but do not appear to actually contribute to the aging index itself.

### Phenotype of aging cells

We next attempted to determine whether epigenetic aging is associated with defined changes in expression phenotype as measured by single-cell RNA-seq. To this end, we analyzed gene expression data from the same individual cells where DNA methylation was measured, focusing first on mouse T cells. Using linear regression models, we correlated the expression of each gene with age (10, 36, 77, or 100 weeks) and with polycomb CpG island methylation index, thus quantifying the effect of time and methylation on the expression levels of each gene, as well as their statistical significance (Supplementary Table 3). Interestingly, many of the genes whose expression in T cells is mostly correlated with polycomb CpG island methylation are associated with defined functional gene families that were already known to be correlated with chronological aging processes^22^. This includes gene sequences involved in protein repair, cell cycle progression, flexibility of sensory perception and basic cell structure integrity (Table 1).

**Table 1.** Single-cell gene expression and DNA methylation in T cells: genes associated with polycomb methylation and chronological age. The table summarizes genes identified from single-cell RNA-seq regression analysis of T cells, modeling expression as a function of biological (methylation at polycomb islands) and chronological (age) measures. Genes are categorized into functional families: protein turnover, cell integrity, sensing, and cell cycle/differentiation, specifying for each gene the standardized regression coefficient of methylation and its adj. p-value, as well as the significance of chronological age coefficient. Known aging-related phenotypes are indicated (e.g., Cancer, Alzheimer’s disease (AD), Parkinson’s disease (PD), immune dysfunction (ID), muscle dysfunction (MD), cardiovascular diseases (CD)). (*) See Methods References.

| Gene family | Gene | Methylation coefficient | Methylation p-val (adj.) | Age p-val (adj.) | Aging phenotype |
| --- | --- | --- | --- | --- | --- |
| Prot. Turnover | Rps29 | -0.37 | <10<sup>-4</sup> | 0.07 | AD<sup>52</sup> |
|  | Rpl35 | -0.30 | <10<sup>-2</sup> | 1 | Cancer<sup>53</sup>, Stroke<sup>54</sup> |
|  | Rnf166 | +0.22 | 0.026 | 0.96 | PD<sup>55</sup> |
| Cell Integrity | Lgals1 | +0.35 | <10<sup>-4</sup> | 0.04 | Cancer<sup>56</sup>, ID<sup>57</sup> |
|  | Actg1 | +0.28 | <10<sup>-2</sup> | 0.5 | Cataract<sup>58</sup>, Hearing<sup>59</sup> |
|  | Ahnak | +0.34 | <10<sup>-4</sup> | 0.03 | NMD<sup>60</sup> |
|  | Vim | +0.26 | <10<sup>-2</sup> | 0.5 | Cancer<sup>61,62</sup>, Cataract<sup>63,64</sup> |
| Sensing | Ccl5 | +0.39 | <10<sup>-6</sup> | <10<sup>-2</sup> | Cancer<sup>65</sup>, AD<sup>66</sup> |
|  | Cd8a | +0.31 | <10<sup>-3</sup> | 0.7 | ID<sup>67,68</sup> |
|  | Nkg7 | +0.25 | <10<sup>-2</sup> | 0.02 | ID<sup>69</sup> |
|  | Cd48 | +0.24 | 0.014 | 0.4 | CD<sup>70</sup> |
| Cell Cycle/Diff. | S100a6 | +0.38 | <10<sup>-6</sup> | <10<sup>-2</sup> | Cancer<sup>71</sup>, AD<sup>72</sup> |
|  | Erdr1 | +0.22 | 0.036 | 1 | ID<sup>73</sup> |
|  | Cdc37 | +0.24 | 0.014 | 0.9 | Cancer<sup>74</sup>, AD<sup>75</sup> |

The most prominent of these genes code for proteins that make up the structural basis for the large (RPL) and small (RPS) ribosomal subunits, which are negatively correlated with aging, consistent with previous findings from bulk tissue samples^23,24^. Our RNA analysis actually revealed over 30 individual ribosomal protein genes that are down regulated as a function of CpG methylation aging (see examples in Table 1). Prominent genes found to be upregulated include those involved in the immune response, cell mobility, the extracellular matrix and cell proliferation, all basic molecular systems that are known to be correlated with the aging phenotype. Indeed, in many cases changes in these individual genes have been shown to be associated with outright characteristics of physiological aging, such as neurodegeneration, hearing loss, cataracts and muscle weakness. It should be noted that in all these examples, regulation is statistically better correlated with epigenetic aging as opposed to chronological age (Supplementary Fig. 3), consistent with the hypothesis that aging is largely a single-cell phenomenon. Indeed, for many specific genes whose association with epigenetic aging was highly significant. In contrast, the corresponding p-value for correlation with chronological aging was found to be insignificant (Table 1).

In order to obtain further verification of these age-related phenotypes, we subjected the same scRNA-seq data to Gene Set Enrichment Analysis (GSEA), a method employed to identify functional classes of genes^25,26^. For this we sorted the genes based on their correlation with either age or methylation and used the GSEA enrichment score to examine if the top or bottom genes are significantly enriched for various gene sets (Methods). Indeed, this computational approach revealed a relatively large number of *a priori* defined gene sets that show statistically significant concordant differences as a function of polycomb CpG island DNA methylation as opposed to chronological age per se, even though the selection of these gene sets was completely unbiased (Supplementary Table 1 and Supplementary Fig. 2). Microanalysis of these sets revealed many of the same age-related genes detected individually (Table 1) from the single-cell RNA library in the gene-specific approach (e.g. RPS, RPL, CD8, Vimentin, Ahnak, etc.), as well as additional age-related changes in RNA that were not previously found by single-cell RNA-seq analysis because they did not pass the statistical threshold. It should be noted that, in general, our ability to identify an aging phenotype was greatly enhanced by employing single-cell regression analysis. This is because the changes in expression observed in individual single cells actually represent only a small fraction of the bulk RNA, making it harder to detect in extracts from the overall cell population.

As noted above, B lymphocytes appear to undergo age-related polycomb CpG methylation at a slower rate than T lymphocytes. To better understand the possible role of this methylation index in B cells, we repeated the above single-cell approach and systematically analyzed how expression changes as a function of methylation and chronological aging (Table 2 and Supplementary Fig. 3). Once again, we observed a strong correlation with gene functions that are known to be affected with advancing age. As opposed to T cells, however, this relationship was much better correlated to chronological age, with the CpG methylation index being less significant.

**Table 2.** Single-cell gene expression and DNA methylation in B cells: genes associated with polycomb methylation and chronological age. Same as Table 1. Here, genes are sorted by age contribution. (*) See Methods References.

| Gene family | Gene | Age coefficient | Methylation p-val (adj.) | Age p-val (adj.) | Aging phenotype |
| --- | --- | --- | --- | --- | --- |
| Prot. Turnover | Rpl41 | +0.25 | 0.78 | <10<sup>-6</sup> | Cancer<sup>76</sup> |
|  | Srrm2 | +0.24 | 0.49 | <10<sup>-5</sup> | Cancer<sup>77</sup>, Dementia<sup>78</sup> |
|  | Rps6 | +0.24 | 0.52 | <10<sup>-5</sup> | Cancer<sup>79</sup>, Lifespan modulation<sup>80,81</sup> |
|  | Hspa5 | +0.22 | 0.96 | <10<sup>-4</sup> | AD<sup>82</sup> |
|  | Sel1l3 | -0.15 | 0.61 | <10<sup>-2</sup> | Cancer<sup>83</sup>, AD<sup>84</sup> |
| Cell Integrity | Mdga1 | -0.25 | 0.78 | <10<sup>-6</sup> | AD<sup>85</sup> |
|  | Tuba1c | +0.21 | 0.95 | <10<sup>-4</sup> | Cancer<sup>86</sup> |
|  | Flna | +0.18 | 0.63 | <10<sup>-3</sup> | Cancer<sup>87</sup> |
|  | Gna13 | +0.19 | 0.39 | <10<sup>-3</sup> | Vascular health<sup>88</sup> |
|  | Ezrin | +0.16 | 0.99 | <10<sup>-2</sup> | Cancer<sup>89</sup>, D<sup>90</sup> |
| Mitochondria | mt-Rnr1 | -0.33 | 0.89 | <10<sup>-11</sup> | Hearing<sup>91</sup> |
| Sensing | Txnip | +0.23 | 0.78 | <10<sup>-5</sup> | Metabolic disorders<sup>92</sup> |
|  | Tsc22d3 | +0.23 | 0.58 | <10<sup>-5</sup> | Cancer, ID<sup>93</sup> |
|  | Cd80 | -0.19 | 0.024 | <10<sup>-3</sup> | ID<sup>94</sup> |
|  | Snx5 | +0.18 | 0.83 | <10<sup>-3</sup> | Cancer<sup>95</sup>, ID<sup>96</sup> |
|  | Cacnb4 | -0.19 | 0.97 | <10<sup>-3</sup> | AD, Memory<sup>97</sup> |
|  | Cxcr4 | +0.16 | 0.50 | <10<sup>-2</sup> | Cancer, ID<sup>98</sup> |
|  | Syk | +0.17 | 0.86 | <10<sup>-2</sup> | Cancer, Arthritis, ID<sup>99</sup> |
| Cell Cycle/Diff. | Tpt1 | +0.18 | 0.95 | <10<sup>-3</sup> | Cancer<sup>100</sup>, ID<sup>101</sup> |
|  | Ets1 | +0.16 | 0.76 | <10<sup>-2</sup> | Cancer<sup>102</sup>, ID<sup>103</sup> |
|  | S100a11 | +0.18 | 0.93 | <10<sup>-3</sup> | ID<sup>104</sup> |
|  | Ier2 | +0.17 | 0.50 | <10<sup>-3</sup> | Cancer<sup>105</sup>, ID<sup>106</sup> |

It should be noted that in addition to T and B cells, functional enrichment analysis strongly suggests an association between polycomb methylation and aging in all cell types (Supplementary Fig. 2). Thus, use of the Enrichr metaserver revealed that genes with significant polycomb methylation coefficients (151 genes with adjusted p-value < 0.05) were significantly enriched (adjusted p-value < 1e-13) in the “Aging Perturbations from GEO up” gene set, containing the top 100 genes in expression signatures comparing young versus old human or mouse tissues, based on chronological age.

Taken together these phenotypic analyses strongly suggest that there are actually two parallel epigenetic aging processes - one that takes place in most cells at a constant and relatively slow rate, combined with a second, more rapid process, which takes place in a more stochastic and independent manner in individual cells. In the case of B cells, the degree of individual single-cell CpG island methylation is relatively minor compared to that taking place in the entire population and, as a result, methylation aging appears to be better correlated with chronological ages.

### Phenotypic aging in single hair cells

While the above analysis using scRNA-seq data clearly suggests that our polycomb CpG island methylation index serves as an excellent clock of biological aging even independent of chronological time, we still sought to determine whether this timing mechanism is also associated with more clear-cut visible characteristics of aging. One of the most obvious signs of human aging is the phenomenon of progressive hair “greying”. Interestingly, a close examination of individual hairs in older individuals shows that this is actually a modular process, with the amount of visual greying not being determined by a uniform degree of whitening in all hair shafts, but rather by the number of individual binary hairs that no longer contain any pigment. This phenomenon itself already suggests that hair aging is not a uniform process, but rather may occur at the single-cell level^27^.

In order to validate that this presumed single-cell aging process is indeed associated with epigenetic aging as measured by the polycomb CpG island methylation index, we plucked individual black or white hairs from a 53-year-old subject (Fig. 5). DNA was extracted from the single cells located at the root of the hair and subjected to scDNA methylation analysis as described (Methods) in order to calculate the aging index. While the majority of black hairs exhibited relatively low methylation levels, most white hairs were found to have a relatively advanced methylation age (Fig. 5). Notably, this differential behavior was observed despite these hairs being taken from the exact same individual at a fixed chronological age. This experiment serves as a poignant validation that biological aging may occur as a single-cell phenomenon, perhaps driven by polycomb CpG island DNA methylation.

**Figure 5:**
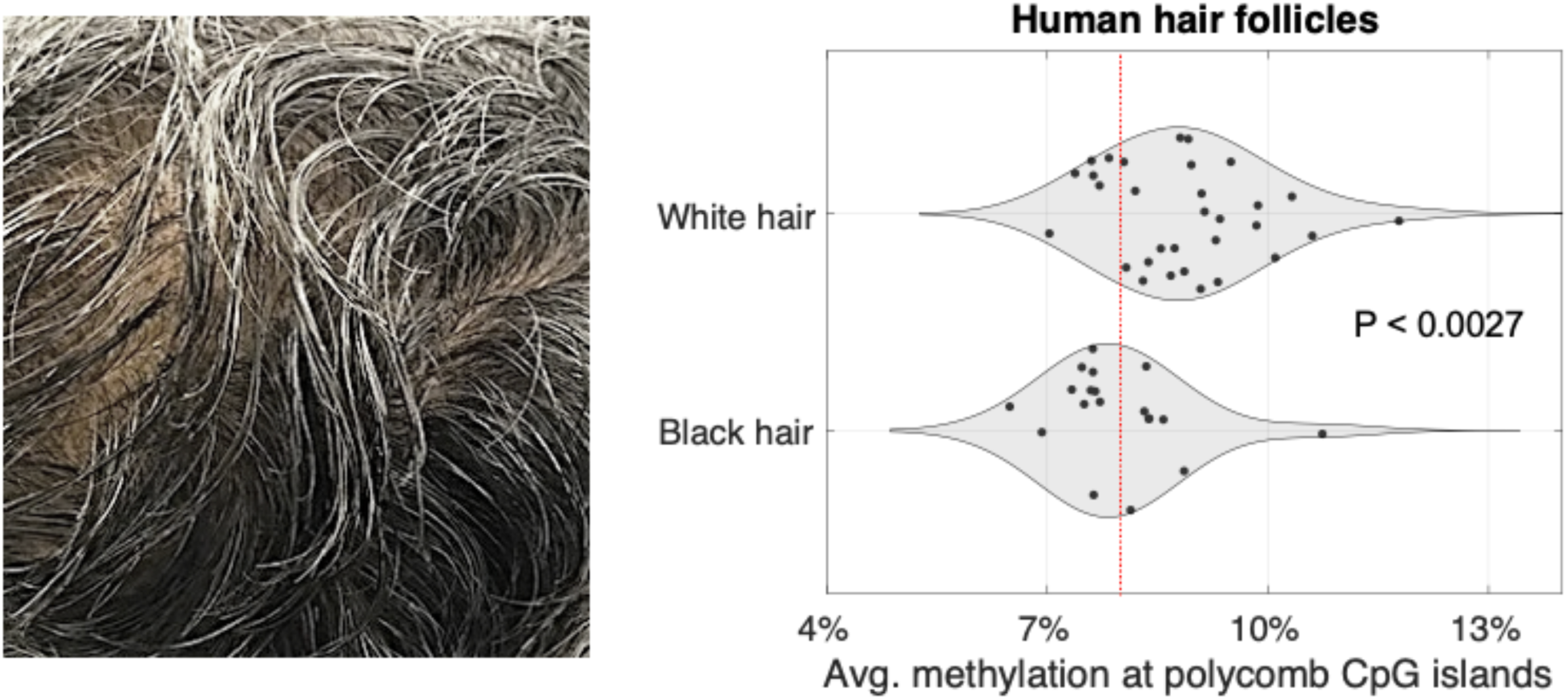
Polycomb CpG island methylation levels in white and black hairs from a 53-year-old individual. Left: an example of our subject’s scalp showing a mix of black and white hairs, highlighting the modular nature of the greying process, where individual hairs either retain pigment (black) or lose it completely (white). Right: polycomb CpG island methylation indices of individual black (n=19) and white (n=31) hairs are presented. White hairs exhibited significantly higher methylation levels compared to black hairs, despite being from the same individual at the same chronological age. The red line emphasizes that most black hair samples (11/19 = 57%) exhibit methylation values below 8%, compared to only 6/31 (19%) white hair samples (p≤0.0027, Kolmogorov-Smirnov test).

## Discussion

The first step in establishing the DNA methylation pattern of all cells in the body occurs very early in development at about the time of implantation, when almost the entire genome undergoes de novo methylation, while CpG islands are protected from this process. This general landscape is then maintained throughout the lifetime of the organism in all cells. Nonetheless, one particular class of CpG islands, those that are bound by the polycomb complex, appear to undergo slow accumulative de novo DNA methylation as a function of age. Indeed, a number of different studies in both human and mouse have demonstrated that the level of polycomb CpG island methylation represents a reliable indicator of chronological age^5,11^ (Supplementary Fig. 1).

### Single-cell aging

In this study, we have attempted to characterize how this process takes place at the topological level. Do all cells in the population undergo methylation aging equally at the same rate, thus constituting a single homogeneous set? Or alternatively, perhaps each individual cell becomes de novo methylated at its own rate; some progressing rapidly and others only very slowly, with the average polycomb CpG island value being proportional to the age of the animal. Our experiments clearly demonstrate that this second model is correct, as many individual cells appear to undergo this methylation aging process at an accelerated rate, independently of the bulk population.

Measuring the methylation age of single cells presented a serious technological challenge, mainly because of sequencing depth limitations and the relative scarcity of CpG residues in the genome. This would normally require carrying out bisulfite sequencing at enormous depth to detect as many polycomb CpG islands as possible. We have overcome this problem by adopting a new methodological approach^11^ whereby all polycomb CpG islands are considered as a single sequence compartment whose average methylation level represents an accurate and reliable measure of age^5^. Using this analytical technique, we examined the single-cell methylation age distribution of many different cell types in the mouse. In every case, the results indicated that while most cells in each population undergo age-dependent methylation at a uniformly slow rate, there are also outliers that show rapid progression of DNA methylation in a manner that deviates from the expected normal distribution. Similar results were obtained for human tissues, as well, in keeping with the concept that polycomb CpG island de novo methylation is a universal marker of aging.

### Age-related demethylation

In addition to polycomb CpG island de novo methylation, aging is also accompanied by the progressive demethylation of many sequences associated with the nuclear lamina (LAD), a large genomic domain that is characterized by replication in late S-phase. Like the phenomenon of de novo methylation, this process appears to be universal in that it occurs in a large variety of different cell types, each at its own rate^20,28^. Using the same domain-wide approach as that for polycomb islands, we were then able to accurately measure the degree of age-related demethylation in single cells. This enabled us to determine whether these two age-related changes in DNA methylation (de novo methylation at polycomb CpG islands and loss of methylation at partially methylated domains) take place in a coordinated manner.

Our studies indicate that cell types which are not known to replicate rapidly do not undergo any appreciable demethylation as a function of age. Cell types with a shorter half-life, however, do show age-related demethylation and this can be seen even at the single-cell level, clearly indicating that these two processes must be mechanistically linked. The exact nature of this connection is not known, but one distinct possibility is that increased polycomb CpG island de novo methylation partially prevents cell differentiation, thus sustaining the replicative state and allowing passive demethylation at LADs^14^.

### Modeling the kinetics of aging

The two modes of epigenetic aging suggest a simple two-state kinetic model for cellular aging. The trajectory of every cell begins in the “young” state, where polycomb CpG island methylation is initially low and subsequently increases at a very low rate. Then, some cells could stochastically and possibly irreversibly switch into a second, “old” state, where polycomb methylation is gained at an accelerated rate. The behavior of different cell types could be accurately explained by different parametrization of this model. Slow-proliferating cells have a near-zero switching probability, resulting in a unimodal distribution, with a slow gain of polycomb CpG island methylation (Fig. 2b). More rapid proliferating cells, on the other hand, randomly switch at different times to the second state, resulting in a bimodal distribution (Fig. 6). This model explains the age-dependent increase of a sub-population of hyper-methylated cells that cannot be explained by cell-cell variability^10^ and provides an intuitive model for the “epigenetic age” of each individual cell (Fig. 6).

**Figure 6:**
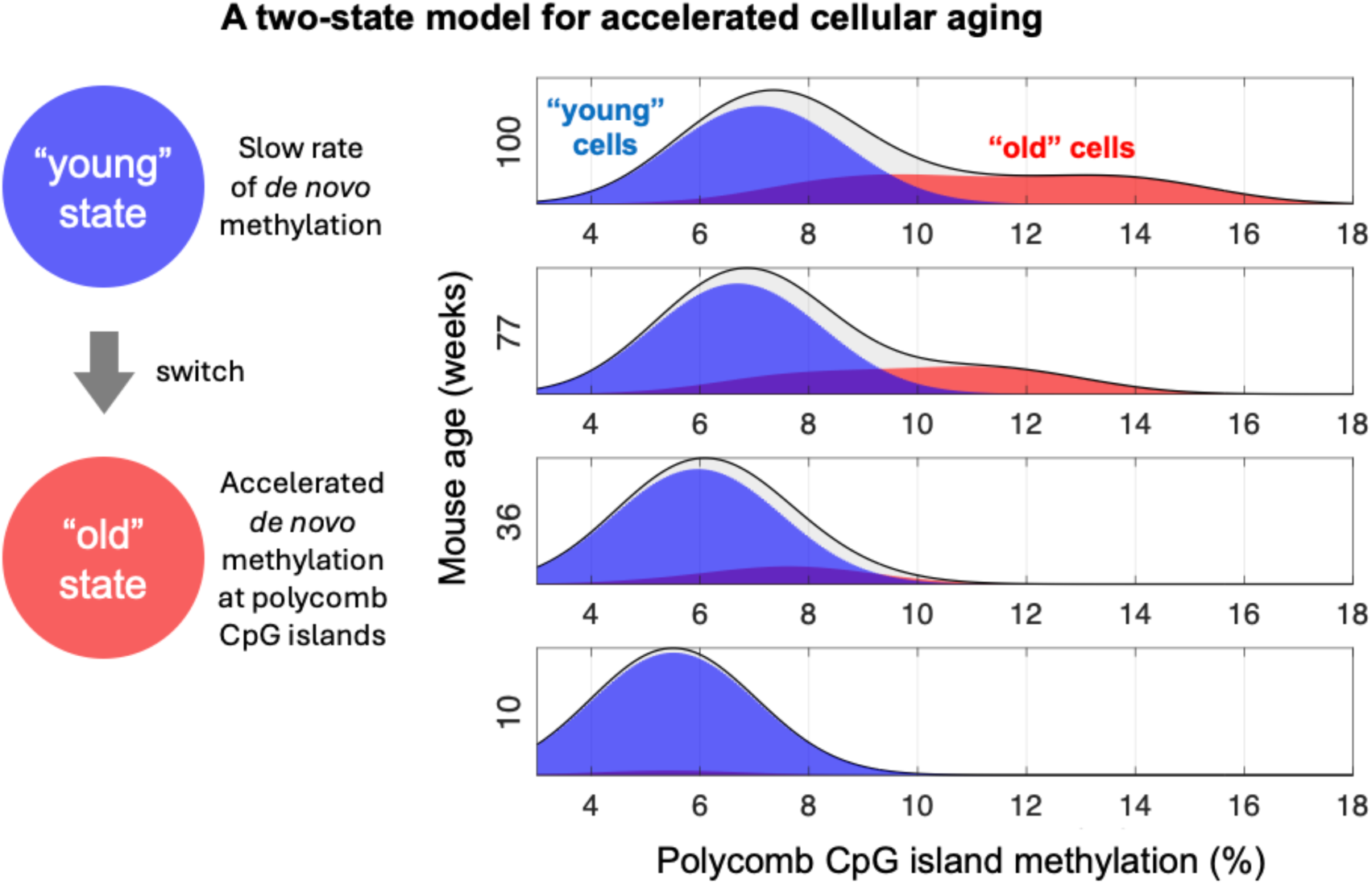
Stochastic Monte Carlo simulation of single-cell polycomb CpG island methylation using a two-state model. The kinetics of single-cell polycomb CpG island methylation is modelled using a simple two-state model (left), composed of a “young” state, by which methylation is gained stochastically at a constant rate. Cells can then stochastically and irreversibly switch to a second state, where the de novo DNA methylation rate is accelerated. Monte Carlo simulations (right) show a gradual age-related increase in blood cells in the “old” state (red), as well as a shift from unimodal to bimodal distribution, with increased cell-to-cell variability in older mice (Fig. 2a, b).

### Phenotype of aging cells

In addition to the direct effect of CpG island de novo methylation on developmental gene promoters, which serves to prevent differentiation and thereby allow continued proliferation^14,29^, these aging cells also appear to acquire their own phenotypic expression profile characterized by the up- or down-regulation of many different gene pathways.

Indeed, a single-cell analysis of DNA methylation and gene expression, obtained from the same individual cells, clearly showed that the polycomb methylation index seems to provide a good reflection of properties associated with biological aging. Using two different analytical approaches, we were able to pinpoint key genes and gene pathways that undergo altered regulation as a function of age-related methylation. This includes genes already known to be associated with functional symptoms typical of old age, and many that were previously undetected; some involved in neurodegeneration, hearing loss, hair growth and the formation of cataracts, as well as those that can affect cell structure, mobility and plasticity^22^. It turned out that our single-cell approach has greatly improved resolution of the aging phenotype. This is because the most significant changes are mainly concentrated in a small number of rapidly aging individual cells, thus overcoming greatly diluted transcription levels in bulk tissue.

Even though the expression pattern of all these genes is strongly correlated with our defined DNA methylation aging index, very few of these genes (<5%) actually have polycomb CpG island promoters; nor are they directly linked to nearby CpG island enhancers. This strongly suggests that while the main driver may be programmed island methylation, generation of the aging phenotype itself is shaped by additional indirect downstream changes in transcription.

These observations may also help in understanding the connection between epigenetic aging and cancer. It is well known that cancer incidence increases with age. It was previously demonstrated that the lifetime cancer risk for each cell type is widely variable, with some tissues (e.g. colon, breast) having a high incidence, while others (e.g. nervous tissues) are very low^30^. Strikingly, these risks are proportional to and can be predicted by the same polycomb CpG island methylation index used here, suggesting that the incidence of tumor formation may actually be a result of the rate of epigenetic aging^5^. The demonstration that methylation aging itself is a single-cell phenomenon clearly provides a basic explanation for why tumors are of monoclonal origin and thus strongly supports a role of epigenetics in cancer. Similarly, many physiological and pathological processes associated with aging, including cataracts, AD, PD and arthritis may originate in individual cells^31–36^.

In addition to the molecular phenotype associated with polycomb CpG island de novo methylation, we showed that this marker is also predictive of the physiological signs of aging. One of the most obvious characteristics of aging in humans is the appearance of “gray” hair, a phenomenon that is generated by the conversion of individual hair follicles from producing their original color to one that has lost this pigment thus sprouting white hair shafts^27^. Our paper now provides a definitive confirmation of this single-cell concept by directly demonstrating that the white hairs indeed have an epigenetically-older age distribution than those that still produce pigment, even when the hairs are taken from the same individual at a fixed chronological age.

### Cause and effect

A major question facing research in this field is whether changes in CpG methylation play a causal role in aging or are simply a consequence of this process. Since most aging methylation indices are derived empirically by following chronological changes in methylation, the sites of methylation change may occur either before or after the phenotypic signs of old age. In contrast, we have demonstrated that polycomb CpG de novo methylation is a programmed process built into the genome, independent of gene expression. Indeed, many tissue- or developmental-specific polycomb-gene promoters become methylated even though those particular genes are not expressed in that cell type, suggesting that expression itself does not play a role in the methylation process^37^.

One of the key characteristics of aging is the increased incidence of cancer. In this case, there are already good reasons to believe that polycomb CpG island de novo methylation, specifically, may indeed be a contributing factor. Not only can this methylation be observed in normal cells prior to their tumorigenic transformation, but the degree of this epigenetic change in different cell types is actually predictive of the lifetime risk of cancer for each tissue^5^. Thus, while the level of polycomb CpG island methylation is relatively elevated in normal intestine, a tissue associated with a high lifetime risk of developing cancer, the amount of methylation in various regions of the normal brain is very low, in keeping with a much lower risk of developing tumors.

We have previously suggested that polycomb CpG island de novo methylation plays a role in tumor formation by inhibiting differentiation^14^. Polycomb is a complex made up of multiple proteins, one of which, Ezh2, is responsible for methylation on lysine 27 of histone H3, thus causing local heterochromatinization that serves to repress underlying gene expression. Its biological role is mainly to silence genes involved in development and differentiation from early in embryogenesis until they are then activated at the appropriate time and position in the body, simply by removing the polycomb complex. Since the promoters of these genes are CpG islands, DNA methylation does not normally play a role in this repression scheme. Like other histone methylases, however, Ezh2 has been shown to have a binding domain that can recruit DNA methylases^15,38^, and as a result, these gene promoters are prone to undergo a slow process of de novo methylation. In combination with the built-in methylation maintenance mechanism, this leads to an accumulation of DNA methylation as a function of aging. This de novo methylation most probably does not directly contribute much to the repression of these genes, since their transcription is already inhibited by virtue of the polycomb complex itself.

Studies on intestinal cancer in the mouse shed light on how DNA methylation may play a critical role in tumor formation. In this system, stem cells located in the crypts generate a continuous supply of epithelial cells that coat the intestinal lumen. After dividing, the daughter cells undergo proliferation as they climb up the crypt, but at one point undergo differentiation to epithelium, thus turning off their proliferative capacity^39,40^. Many of the genes involved in this process are controlled by polycomb^7^. Normally, they would be activated by removal of this complex, but if these cells have already undergone considerable aging through the accumulation of CpG island methylation, they can no longer be turned on, even if the polycomb complex is released. In the absence of differentiation, these cells will then continue to proliferate, thus contributing a major component of tumorigenesis^14^.

Taken together, these observations clearly suggest that the generation of tumors can be a direct result of methylation aging. According to this concept, cancer has a single-cell origin because aging itself takes place at the level of single cells. We propose that many of the non-tumor characteristics associated with aging can also develop in individual cells as a result of decreased plasticity in the differentiation process. This, for example, could explain the many age-associated changes that occur in the immune and other sensing systems^41,42^, which slowly lose their flexibility to properly activate key differentiation genes in response to changes in environment.

## Methods

### Single-crypt isolation and whole-genome methylation library preparation

Small intestine crypts were isolated from two wildtype B6 mice, aged 22 and 28 weeks, as previously described^107^. Briefly, the proximal half of the small intestine was isolated, opened longitudinally, and washed with ice-cold PBS to clear most of the luminal contents. The intestine was then cut into small fragments (0.5-1 cm) and washed again 2-3 times. Intestinal fragments were incubated in 2 mM EDTA in DPBS at 4°C for 30-45 minutes with gentle shaking. After incubation, the tube was shaken vigorously to dislodge crypts. Suspension was filtered through a 70 µm cell strainer to remove debris. Crypts were resuspended in PBS, and under a light microscope, individual crypts were carefully isolated using a 10 µL pipette. Each single isolated crypt was then transferred to a small PCR tube and stored at -80°C for subsequent DNA extraction. For DNA extraction, each crypt was resuspended in 20 µL of lysis buffer (20 mM Tris, 20 mM EDTA, 20 mM KCl, and 0.3% Triton X in double-distilled water), with 80 µg of proteinase K added per sample. The samples were incubated at 50°C for 3 hours, followed by 85°C for 10 minutes, and then cooled on ice or stored at - 20°C. Subsequently, sonication was performed (see below), and whole-genome methylation libraries were prepared using the NEBNext Enzymatic Methyl-seq Kit (E7120) according to the manufacturer’s instructions. Sequencing was carried out on the Illumina NovaSeq platform with paired-end 150 bp reads, covering approximately 1× of the genome.

### Single-hair isolation and WGBS-library preparation

Individual black or white hairs were plucked from a 53-year-old subject by a dermatologist. The hairs were grouped into black and white categories and immediately subjected to DNA extraction with the QIAGEN DNeasy Blood & Tissue Kit (Cat. No. 69504) following a user-developed protocol^108^. In brief, hair samples were cut into 0.1-1 cm piece from the base (root) and transferred to a 1.5 mL microcentrifuge tube containing 300 µL Buffer ATL, 20 µL Proteinase K, and 20 µL of 1 M DTT. The samples were mixed by pulse vortexing for 10 seconds and incubated at 56°C until completely lysed (at least 1 hour or overnight). After lysis, the DNA samples were vortexed precipitated with ethanol according to the protocol. The extracted DNA samples were stored at -20°C until sonication was performed (see below). Bisulfite conversion was carried out using the EZ DNA Methylation-GOLD Kit (Cat. No. D5005). Whole-genome bisulfite sequencing (WGBS) libraries were generated using the xGen Methyl-seq Library Prep Kit (10009824, IDT). Sequencing was performed on the Illumina NovaSeq platform with paired-end reads of 150 bp from each side, as described above. DNA shearing was performed using the Bioruptor Pico (Diagenode). Between 10–12 shearing cycles were performed using the 30’’/30’’ Easy mode program. Each 50 µL sample (in TE buffer with low EDTA) was placed in 0.2 mL microtubes for the Bioruptor Pico (C30010020). The samples were mixed and spinned down after every three cycles to ensure uniform shearing.

### WGBS data processing

FastQ files were processed with the nf-core/methylseq pipeline using the BISCUIT aligner as previously described^109,110^. Briefly, the quality of fastQ files was assessed using FastQC (v0.11.9). TrimGalore (v0.6.5) was used to trim adapters from all samples with the parameters: “clip_r1=10 clip_r2=15 three_prime_clip_r1=10 three_prime_clip_r2=10”. We aligned the trimmed samples to the mm10 genome assembly using the biscuit aligner (0.3.16.20200420) with “-b 1” for directional library. Duplicates were marked using samblaster (v0.1.24) with “--addMateTags --excludeDups”. Then, BAM files were sorted and indexed using SAMtools (v1.9). The sorted and indexed BAMs were given as an input to biscuit pileup to generate VCF files. Bed files were generated from VCF using biscuit vcf2bed with “-k 1 -t cg” arguments. Additionally, Bed files were filtered to include only CpGs in the solo-WCGW context^20^ using BEDtools 2.29.1. Both regular and solo-WCGW BED files were further filtered to remove CpGs in blacklisted genomic regions^111^. A custom SNP-filtering approach was implemented using BCFtools (v1.9), and BISCUIT epiread files were generated by merging paired-end reads. Library quality control was performed based on the number of mapped reads per sample; samples with low read depth (<2M reads) were excluded from downstream analysis. Additional quality control metrics, including insert size distribution, duplication rates, error rates, and non-CpG methylation levels, were systematically evaluated for each sample. Samples exhibiting outlier values in these metrics were removed prior to further analysis. A MultiQC report generated by the nf-core/methylseq pipeline (https://github.com/ekushele/methylseq) was used to aggregate and evaluate all quality control metrics across samples (Supplementary Dataset 3).

### Calculation of average methylation index

Average methylation was calculated at polycomb CpG islands in mouse (n=2,975) and human (n=500) islands^5^ (Supplementary Datasets 1,2). WGBS or EM-seq methylation data were used to identify overlapping reads (bedtools intersect). For each region, the average methylation was calculated as the total number of methylated CpGs divided by the total number of observed CpGs, in this region. Regions with fewer than 10 binary observations were discarded. The methylation index for each individual cell was then calculated by averaging across these estimates. Following these steps, samples with a low proportion of reads mapping to Polycomb islands were additionally excluded from the analysis. The methylation index for each individual cell was then calculated by averaging across these estimates (Supplementary Dataset 4). Similarly, the average methylation at Solo-WCGW sites ([AT]CG[AT], with no flanking CpGs at 35bp) was computed, within the PMD regions of the genome. Actual distributions of polycomb CpG island methylation was also compared to simulated data, sampled from a normal distribution, while maintaining the same size, mean, and standard deviation as the observed data (in each island). To ensure biological relevance, we adjusted the simulated values to remain within the observed range, as methylation at CpG polycomb islands can only increase^5^.

### Single-cell RNA-seq data analysis

#### Data Acquisition and Pre-processing

scRNA-seq data from white blood cells were obtained from GSE225173. Analysis was performed in R (version 4.3.0) using the Seurat package. A Seurat object was created, including only cells with a minimum of 1,500 detected genes and genes expressed in at least three cells. The percentage of mitochondrial gene content was calculated for each cell. Based on quality control (QC) criteria, only cells meeting the following thresholds were retained: Gene count < 4,700, Unique molecular identifier (UMI) count < 850,000 and Mitochondrial gene percentage < 15%. Dimensionality reduction was performed using principal component analysis (PCA), followed by uniform manifold approximation and projection (UMAP) for visualization, using the first 20 principal components, as determined by an elbow plot. Clustering was conducted with a resolution parameter of 0.4.

#### Cell-type annotation and subsetting

Cell types were determined for each individual single cell using canonical marker genes, as previously described^10^. These include T cells (*Cd3e, Cd3d, Cd3g*), B cells (*Cd19, Cd79a, Cd79b*), natural killer (NK) cells, and monocytes. Following QC and cell-type identification, the dataset was subsetted to retain only T cells (n = 301) and B cells (n = 599). Cells lacking average polycomb CpG methylation data were excluded from the analysis.

#### Gene expression and regression analysis

Each cell type subset was analyzed independently following the same workflow: **1.** Normalization was performed using SCTransform (v2 regularization). **2.** Genes not expressed in at least 25% of cells were excluded, leaving 3,431 genes in the T cell subset and 2,725 genes in the B cell subset. **3.** A multivariable linear regression model was fitted for each gene, treating normalized gene expression as the dependent variable (each cell as an independent observation). The model was specified as:

*GeneExpression = β_0_ + β_1_ × ChronologicalAge + β_2_ + PolycombMethylation + ε*

where β₀ is the intercept, β₁ is the coefficient for chronological age (modeled as a continuous variable with values 10, 36, 77, and 100 weeks), β₂ is the coefficient for average polycomb CpG island methylation level, per cell, and ε denotes the error term.

Linear modeling was then performed using lm function from base R stats package. Standardized coefficients were obtained using the lm.beta() function from the QuantPsyc R package. **4.** Multiple hypothesis correction was applied separately to the p-values of each variable using the Benjamini-Hochberg procedure. The same analytical approach was applied to scRNA-seq data from muscle stem cells obtained from GSE121364. The full results are provided in Supplementary Table 3. Volcano plots were generated to visualize the relative influence of polycomb CpG methylation and chronological age on gene expression across all single B and T cells. For each gene, standardized coefficient estimates from the linear regression model were plotted against their corresponding adjusted p-values (−log₁₀ scale), separately for the chronological age (actual age) and biological age (polycomb CpG islands methylation) coefficients. This approach allows visualization of the relative contribution of each variable to gene expression variation.

### Gene Set Enrichment Analysis (GSEA)

GSEA was conducted using the GENI^26^ online service. For ranking, we used the full regression models and ranked genes by their correlation coefficients and its statistical significance, using (−log₁₀ (p-value) × sign (regression coefficient), which was calculated separately for polycomb methylation and for chronological age (Supplementary Tables 1-3). For cross-species comparisons, genes were mapped to their human orthologs using the “orthogene” package and non-conserved genes were omitted.

### Two-state model for DNA methylation in aging

Single-cell shallow whole-genome DNA methylation data^10^, was obtained for 1,132 blood cells from mice aged 10, 36, 77 and 100 weeks. Single-cell RNA-seq data was used to classify each cell, and average DNA methylation level was computed and averaged across polycomb CpG islands, as described above. Cells were grouped into non-proliferating cells (B-cells, Naive CD4 T-cells, and Naive CD8 T-cells) and cells showing high proliferation rates (Effector CD4 T-Cells, Memory CD8 T-cells, Regulatory T-cells, and NK cells). To model the stochastic accumulation of methylation with time, we first proposed a “one-state” model, by which methylation is gained, per cell, at a constant stochastic rate. In a second model, cells can stochastically and irreversibly switch to a second state, where DNA methylation is gained at another rate. We then used Monte Carlo simulations, to optimize the parameters of the two models, to fit the actual single-cell data. For the non-proliferating cells, the probability of state-switching was ∼0.2%/week, with a total of ∼18% of cells reaching the ”older” state by week 100. The proliferating cells show a much higher rate of switching, ∼0.65%/week, with ∼40% of “old” cells at week 77 and nearly 50% at week 100. The average gain of methylation was ∼0.02 percent points per week for the “young” state, and ∼0.1%/week for the “older state”.

### Data

Methylation data for mouse: ESLAM and LSK cells (GSE89545), Muscle stem cells (GSE121436), White blood cells (GSE225173). Methylation data for human: fetal liver, cord blood#2, bone marrow megakaryocytes, common lymphoid progenitors (CLP) and common myeloid progenitors (CMP) (GSE87196/^44^). Cord blood#1 (GSE106957^43^). scRNA-seq data mouse: White blood cells (GSE225172), muscle stem cells (GSE121437).

## Code availability

Relevant code is available at https://github.com/shmuel-ruppo/cellular-aging/.

## Acknowledgments

We thank Prof. Avi-Hai Hovav for his willingness to provide single hair strands, and Ms. Tzippi Jakubowicz for organizing and editing the manuscript for publication. We also acknowledge support from the Howard Jonas Foundation, the Rosetrees Foundation, the Israel Science Foundation, and the Israel Cancer Research Foundation, the Biotechnology and Biological Sciences Research Council, BBSRC and Wellcome Trust. Y.D., T.K. and H.C. are members of the Pamela and Paul Austin Research Center on Aging, at the Hebrew University.

## Author contributions

H.M. designed and conducted the experiments, performed the WGBS data analysis (including newly generated data and publicly available datasets), assisted with the single cell RNA data analysis, interpreted the results, and contributed to manuscript writing and preparation. S.R. performed the single cell RNA data analysis. S.J.C. M.J.B., and F.V.M. contributed to creating the mouse blood and muscle stem cell data. M.H. assisted with designing the single crypt experiments. E.K. and S.O. assisted with WGBS data analyses. O.V. assisted with the single cell RNA data analysis. A.Z. directed the single-hair experiments. S.E. organized the single cell RNA data analysis. Y.D. contributed to manuscript writing and preparation. W.R. contributed mouse single cell data and manuscript writing and preparation. T.K. directed the study, developed a computational model for the kinetics of aging, contributed to data analysis and interpretation, and assisted in manuscript writing and preparation. H.C. directed the study and wrote the manuscript.

## Competing interests

The authors declare no competing interests.

## Supplementary Data

**Supplementary Fig. 1:**
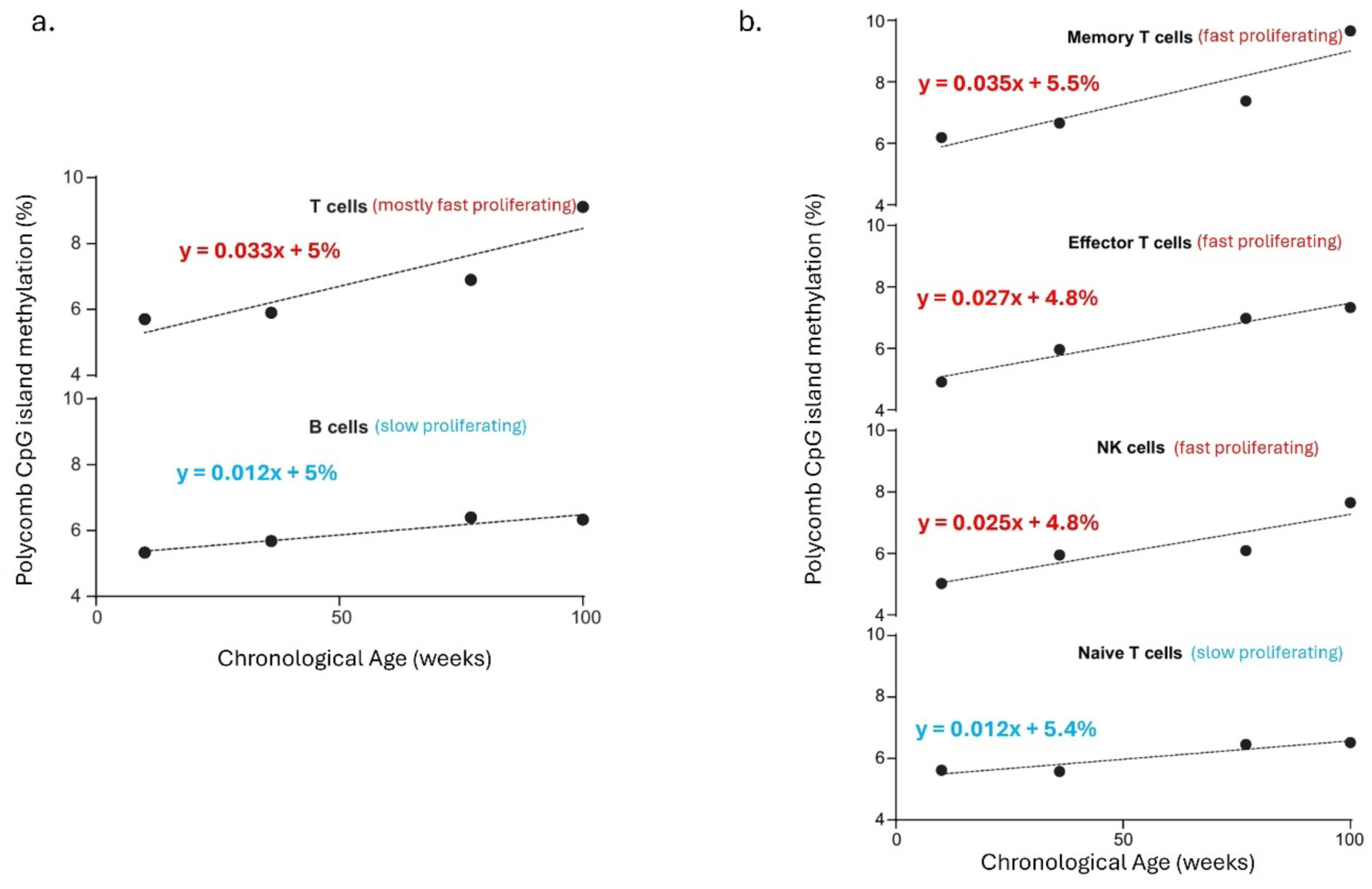
Chronological age and cell proliferation correlates with polycomb CpG island methylation. **(a)** The relationship between polycomb CpG island methylation levels (%) and chronological age (weeks) is shown for T cells and B cells. Each data point represents the average methylation level across all analyzed cells at the respective time point. Linear regression analysis reveals that polycomb CpG island methylation accumulates with age in both cell types, with T cells showing a steeper slope (0.033) compared to B cells (0.012). This indicates that proliferating T cells exhibit a faster rate of methylation accumulation than less proliferative B cells. **(b)** Polycomb CpG island average methylation levels (%) in T cell subtypes across the indicated chronological ages. The rate of methylation accumulation varies among subtypes, with Memory T cells showing the steepest increase (0.0346), followed by Effector T cells (0.027), NK cells (0.025), and Naive T cells (0.012). The highest slope observed in Memory T cells is nearly three times greater than the lowest slope found in Naive T cells. These results demonstrate subtype-specific methylation dynamics, further emphasizing the significant differences in the rate of methylation accumulation between different cell subtypes. Source of data: GSE225173.

**Supplementary Fig. 2:**
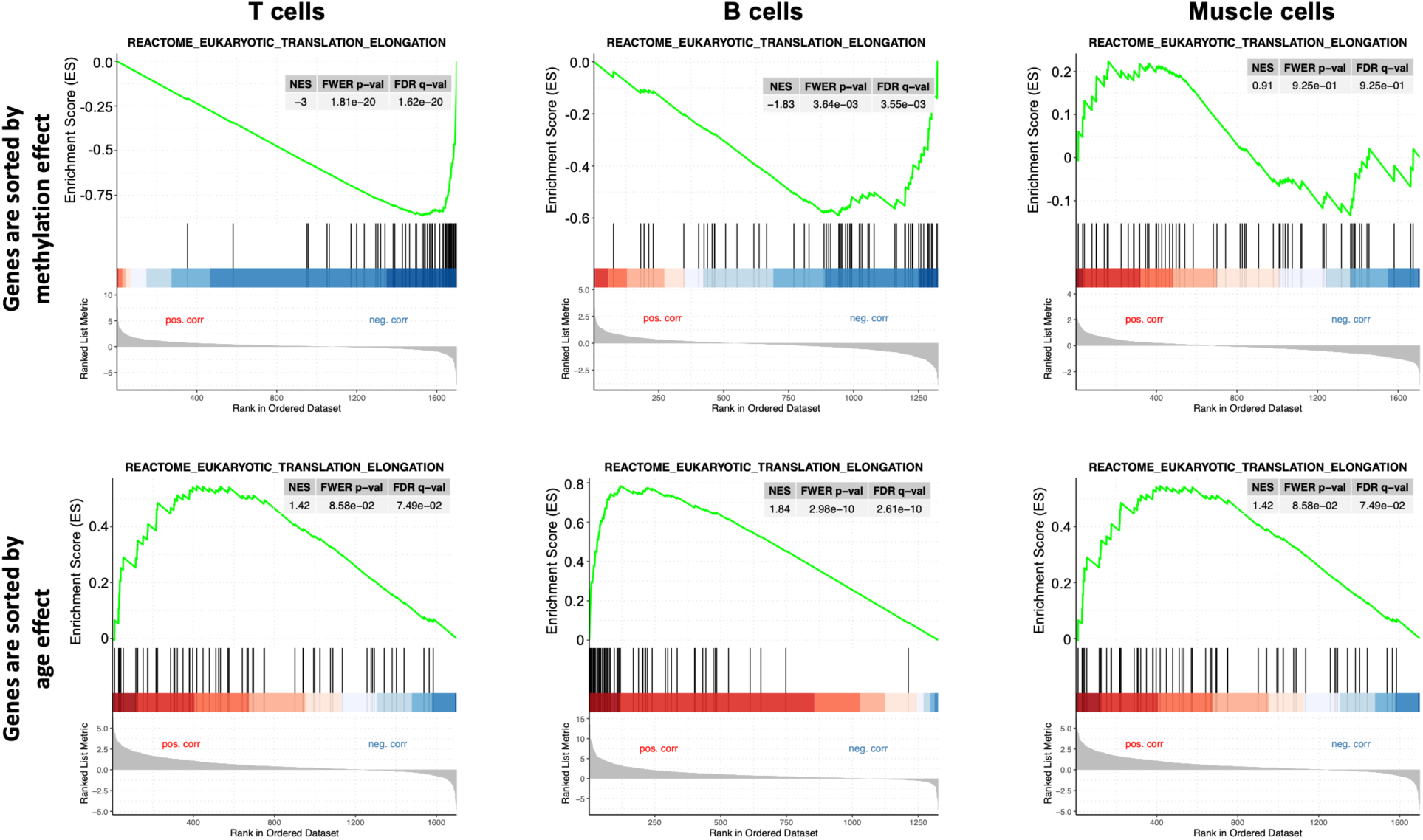
Gene Set Enrichment Analysis (GSEA) Examples. Each panel depicts the GSEA enrichment for the “Translation Elongation” gene set (Reactome Pathway Database) under distinct experimental conditions (Methods). For each condition (T cells, B cells, Muscle cells), genes were sorted based on their methylation (top) or age (bottom) coefficient in a full regression model using single-cell gene expression and DNA methylation data. The green line represents the running Enrichment Score (ES) as pathway-related genes are encountered in the ranked gene list. The vertical bars below highlight the ranked positions of pathway genes, with red indicating upregulated genes (positive correlation) and blue downregulated genes (negative correlation). Statistical values, including normalized enrichment score (NES), family-wise error rate (FWER) p-value, and false discovery rate (FDR) q-value, are shown in each panel. Positive NES values indicate upregulation of the pathway, while negative NES values reflect downregulation.

**Supplementary Fig. 3:**
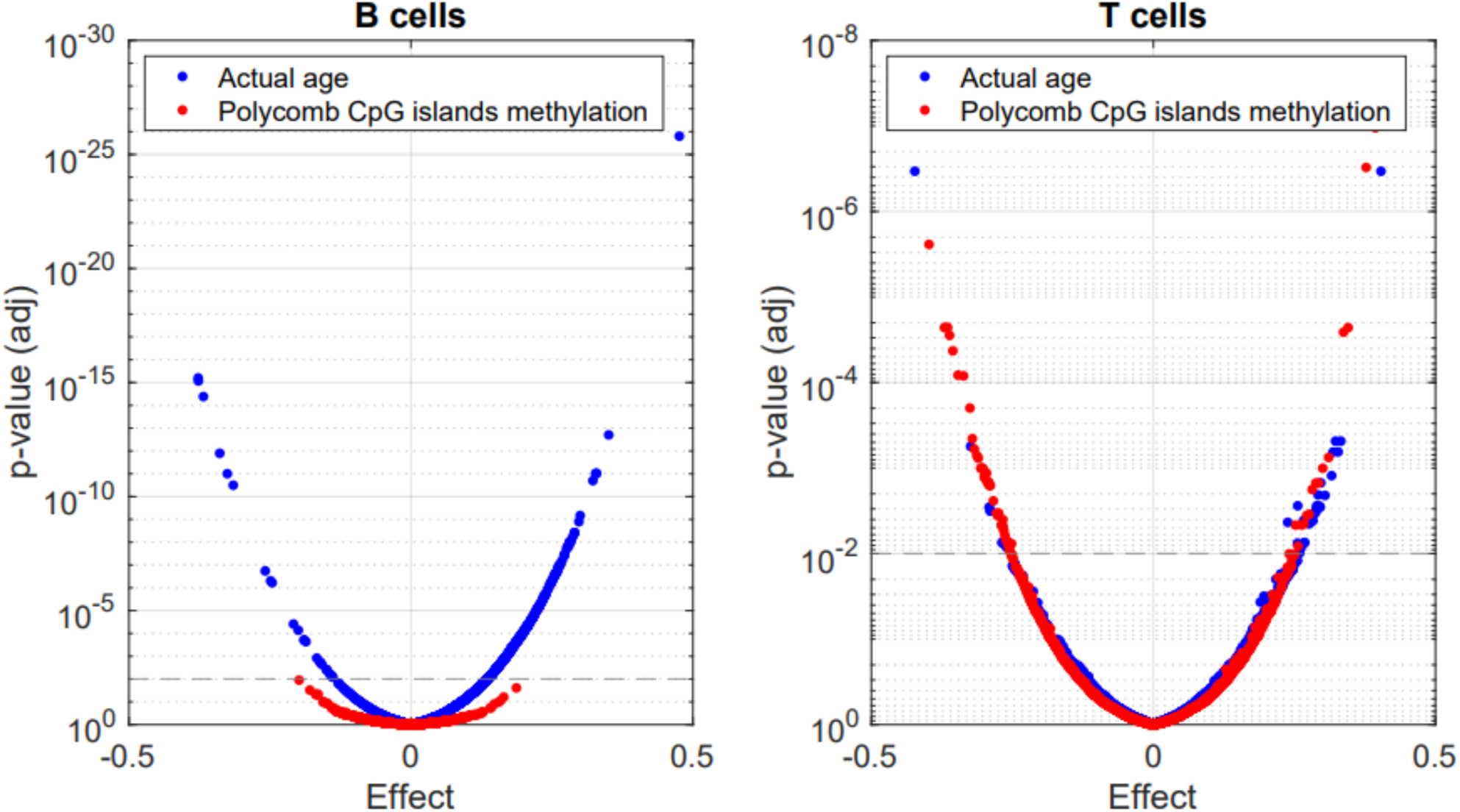
General association between phenotype and age in B and T cells. Volcano plot shows adjusted p-value for the association between both positive and negative changes in gene expression as a function of chronological age (blue) or polycomb CpG island methylation as an index of aging (red) in single B or T cells from mouse. Note that in B cells, in which polycomb CpG island de novo methylation does not increase, expression changes are only significant as a function of chronological age. In T cells one sees significant genes (p<1e-2) for both polycomb and chronological age, consistent with the observation that these cells undergo two types of aging (bulk and single-cell). Nonetheless, at the level of p<1e-3, there are indeed more significant genes (18) when comparing polycomb methylation vs gene expression but only 7 using chronological age.

**Supplementary Table 1:** Gene Set Enrichment Analysis (GSEA) results for T cells. The table summarizes the results of GSEA performed on the scRNA-seq data from T cells, comparing gene expression changes as a function of polycomb CpG island DNA methylation and chronological age. GSEA was conducted using the GENI web server^26^, with genes ranked based on -log10 (p-value) × sign (regression coefficient), calculated separately for polycomb methylation levels and chronological age (see Methods). Gene sets (column 2) are grouped into functional categories (column 1: Gene family), including Protein Turnover, Cell Integrity, Sensing, and Cell Cycle/Differentiation. For each gene set, the table reports results for both polycomb methylation age (biological age) and chronological age, including; Normalized Enrichment Score (NES): A measure of the strength and direction of enrichment and False Discovery Rate (FDR) q-value: Adjusted p-value accounting for multiple comparisons. Gene sets were from Reactome, except *Cytoskeletal fiber* and *Cell-cell junction*, which were from GO Cellular Component.

|  | **GSEA T cells** | Polycomb methylation age |  | Chronological age |  |
| --- | --- | --- | --- | --- | --- |
| Gene family | Gene set | NES | FDR (q-value) | NES | FDR (q-value) |
| Prot. Turnover | Translation | -2.8 | <10<sup>-16</sup> | 1.4 | 0.05 |
|  | RRNA processing | -3.0 | <10<sup>-19</sup> | 1.4 | 0.08 |
| Cell Integrity | Cytoskeletal fiber | 1.8 | 0.001 | 0.5 | 0.8 |
|  | Cell-cell junction | 1.6 | 0.02 | 0.85 | 0.8 |
| Sensing | Influenza infection reactome | -3.0 | <10<sup>-19</sup> | 1.6 | 0.03 |
|  | Infectious disease response | -1.9 | <10<sup>-6</sup> | 1.3 | 0.04 |
|  | Cellular response to stimuli | -2.2 | <10<sup>-10</sup> | 1.4 | 0.03 |
| Cell Cycle/Diff | Nervous system Development | -2.2 | <10<sup>-8</sup> | 1.4 | 0.03 |
|  | Developmental biology | -2.1 | <10<sup>-9</sup> | 1.3 | 0.04 |

**Supplementary Table 2:**
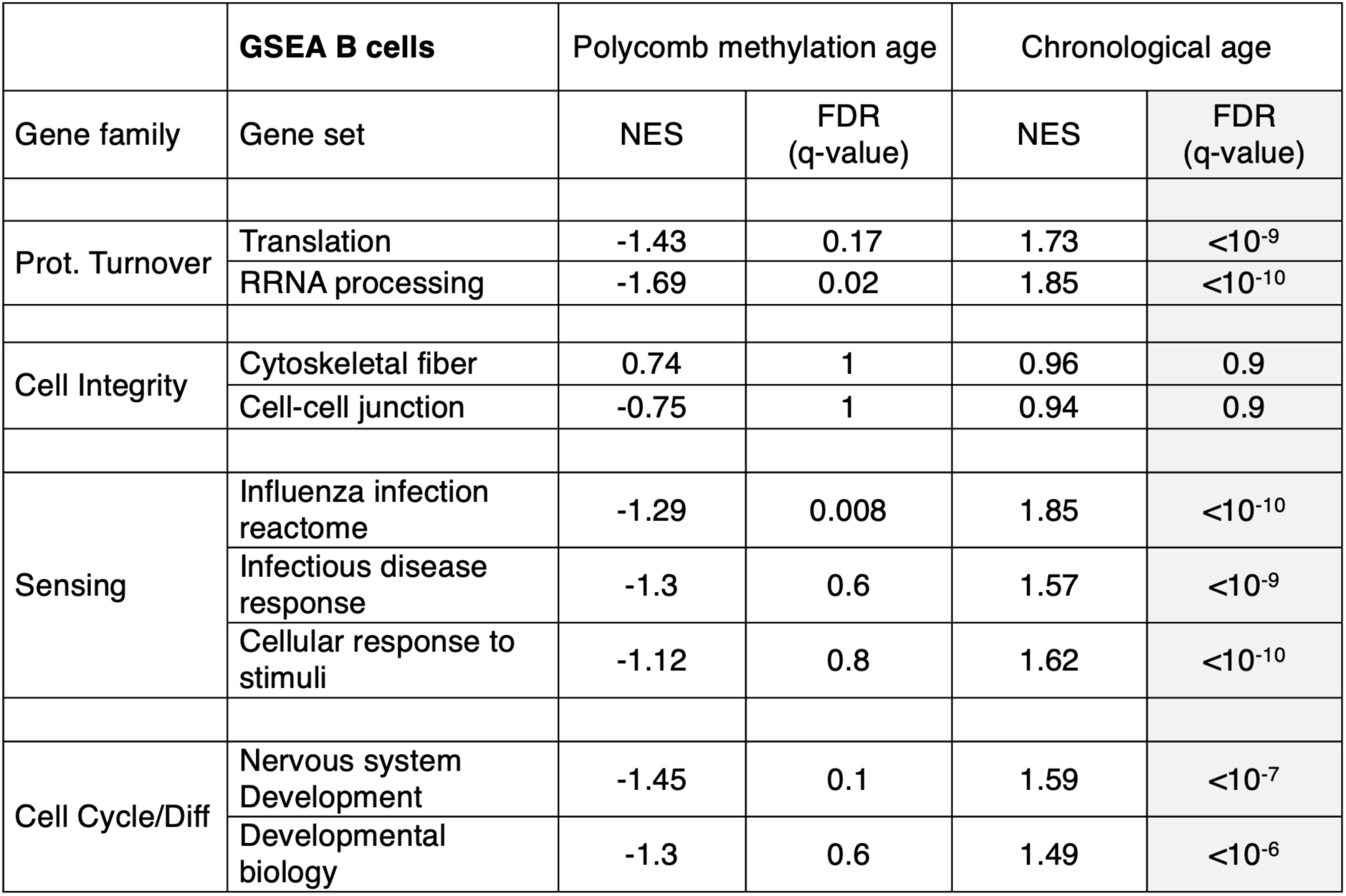
Gene Set Enrichment Analysis (GSEA) results for B cells. The table presents the GSEA results for B cells, following the same analysis approach as described in Supplementary Table 1.

**Supplementary Table 3: Full gene-specific regression models for T and B cells (age and methylation)**

Supplementary Dataset 1: List of 2,975 polycomb CpG islands in mouse (mm10)

Supplementary Dataset 2: List of 500 polycomb CpG islands in human (hg19)

Supplementary Dataset 3: WGBS quality control information.

Supplementary Dataset 4: DNA methylation per island per sample.

## Notes

### Competing Interest Statement

The authors have declared no competing interest.

